# Metabolic contributions of an alphaproteobacterial endosymbiont in the apicomplexan *Cardiosporidium cionae*

**DOI:** 10.1101/2020.10.19.346205

**Authors:** Elizabeth Sage Hunter, Christopher J Paight, Christopher E Lane

## Abstract

Apicomplexa is a diverse protistan phylum composed almost exclusively of metazoan-infecting parasites, including the causative agents of malaria, cryptosporidiosis, and toxoplasmosis. A single apicomplexan genus, *Nephromyces*, was described in 2010 as a mutualist partner to its tunicate host. Here we present genomic and transcriptomic data from the parasitic sister species to this mutualist, *Cardiosporidium cionae,* and its associated bacterial endosymbiont. *Cardiosporidium cionae* and *Nephromyces* both infect tunicate hosts, localize to similar organs within these hosts, and maintain bacterial endosymbionts. Though many other protists are known to harbor bacterial endosymbionts, these associations are completely unknown in Apicomplexa outside of the Nephromycidae clade. Our data indicate that a vertically transmitted **α**-proteobacteria has been retained in each lineage since *Nephromyces* and *Cardiosporidium* diverged. This **α**-proteobacterial endosymbiont has highly reduced metabolic capabilities, but contributes the essential amino acid lysine, and essential cofactor lipoic acid to *C. cionae*. This partnership likely reduces resource competition with the tunicate host. However, our data indicate that the contribution of the single **α**-proteobacterial endosymbiont in *C. cionae* is minimal compared to the three taxa of endosymbionts present in the *Nephromyces* system, and is a potential explanation for the virulence disparity between these lineages.

## Introduction

Apicomplexa includes a multitude of highly virulent pathogenic organisms, such as *Plasmodium falciparum*, *Cryptosporidium parvum*, and *Toxoplasma gondii,* the causative agents of malaria, cryptosporidiosis, and toxoplasmosis, respectively. Malaria claims about half a million human lives annually (Center for Disease Control 2019), *T. gondii* is estimated to infect up to 60% of the human population in much of Europe (Pappas, Roussos, and Falagas 2009), and cryptosporidiosis causes 3-5 million cases of gastrointestinal disease annually in children in Africa and India alone (Sow et al. 2016). These organisms represent major human health concerns, but as a result, our understanding of this phylum is largely based on a small subset of clinically relevant apicomplexans. Every metazoan likely plays host to at least one apicomplexan (Morrison 2009), and this is probably an underestimation, as many species can host multiple apicomplexan species. Apicomplexans have been described in a vast array of vertebrates from avians to marine mammals (Jeurissen et al. 1996; Conrad et al. 2005), and also in cnidarians (Kwong et al. 2019), molluscs (Suja et al. 2016; Dyson, Grahame, and Evennett 1993), arthropods (Criado-Fornelio et al. 2017; Alarcón et al. 2017), and urochordates (Ciancio et al. 2008; Mary Beth Saffo et al. 2010). Their host range is enormous, and their diversity and adaptation to the parasitic lifestyle is unparalleled.

The long history of evolution and adaptation to life within a host has given rise to a series of characteristic genomic losses and the evolution of specialized cellular machinery in apicomplexans (Roos 2005; Janouskovec and Keeling 2016; McFadden and Waller 1997; Soldati, Dubremetz, and Lebrun 2001; Frénal et al. 2017). Specific structural adaptations of these organisms include those for functions related to host infection and persistence; namely a remnant plastid (apicoplast) and apical complex (McFadden and Waller 1997; Soldati, Dubremetz, and Lebrun 2001). Genomic reductions associated with parasitism in apicomplexans include losses in gene families for the biosynthesis of purines, amino acids, sterols, various cofactors, the glyoxylate cycle, endomembrane components, and genes related to motility (Janouskovec and Keeling 2016; Woo et al. 2015). Additionally, apicomplexans also show expansions in gene families related to infection and persistence within host cells (Janouskovec and Keeling 2016). However, the assumption that these genomic signatures are associated with parasitism is based on limited information, since a direct comparison to closely related free-living sister taxa is not possible, and there are no known free-living apicomplexans (Janouskovec and Keeling 2016). However genomic data is available from the photosynthetic Chromerids (Woo et al. 2015), which likely diverged from apicomplexans 600-800 million years ago (Votýpka et al. 2016).

Despite the high pathogenicity and parasitic adaptations of many members, questions have emerged over whether Apicomplexa is an entirely parasitic group. Though this sentiment has long been mentioned in publications (Morrison 2009; Roos 2005; Mathur et al. 2018; Gubbels and Duraisingh 2012; McFadden and Yeh 2017; Woo et al. 2015, Votýpka et al. 2016), the current evidence suggests that the interactions between apicomplexans and their hosts are far more varied than previously recognized. In fact, it is likely that apicomplexans span the full spectrum from parasitism to commensalism, and even mutualism (Rueckert, Betts, and Tsaousis 2019; Kwong et al. 2019; Saffo et al. 2010). However, what defines the boundaries along this continuum of symbiotic association is still a topic of much debate (Leung and Poulin 2008; Johnson and Oelmüller 2009; Ewald 1987). Phylogenetic analysis indicates *Nephromyces* is sister to the haematozoan clade, and closely related to highly virulent genera such as *Plasmodium, Theiliera,* and *Babesia* (Muñoz-Gómez et al. 2019). Thus far, apicomplexan species with variable life strategies have been found in early branching groups, such as the Gregarina and Corallicods. However, the existence of this reportedly mutualistic taxon deep within Apicomplexa, sister to a group of highly virulent blood parasites, suggests the unique biology of Nephromycidae might be responsible for such a shift to a commensal or mutualistic life strategy.

*Cardiosporidium cionae* was originally described in 1907 by Van Gaver and Stephan, who correctly identified it as a novel sporozoan parasite of the invasive tunicate *Ciona intestinalis*. This species wasn’t mentioned again until it was observed by Scippa, Ciancio, and de Vincentiis in 2000, and then formally redescribed by Ciancio et al. in 2008, a full century after its initial discovery. Similar to other haemosporidians such as *Plasmodium*, *C. cionae* is found in the blood of its host. It localizes to the heart and pericardial body, a collection of sloughed off cells that accumulates over the life of the tunicate inside the pericardium (Evans Anderson and Christiaen, 2016). *Ciona intestinalis* is highly invasive; this prolific species has spread globally traveling in the hulls and bilgewater of ships and is now found on every continent except Antarctica. While *C. cionae* infection has only been formally confirmed in The Gulf of Naples, Italy (Ciancio et al. in 2008), and Narragansett Bay, Rhode Island, USA, it likely has a broad range as well. Additionally, TEM data from the redescription of *C. cionae* revealed a bacterial endosymbiont (Ciancio et al. 2008).

The closest relative of *C. cionae*, *Nephromyces,* was first described around the same time in 1888, though its unusual filamentous morphology caused it to be misclassified as a chytrid fungus until 2010 (Saffo et al. 2010). *Nephromyces* is found in the Molgulidae family of tunicates, in a ductless structure of unknown function adjacent to the heart, known as the renal sac. It is thought to be mutualistic due to a near 100% infection prevalence (Saffo et al. 2010) and is capable of utilizing the waste products that the host tunicate sequesters in the renal sac as a source of glycine, pyruvate, and malate (Paight et. al 2019).

*Nephromyces* also houses separate three lineages of bacterial endosymbionts (Paight et al. 2020). Though endosymbiotic associations are commonly found in other protists such as ciliates, diatoms, and amoebas, bacterial endosymbiosis in Apicomplexa is unique to this lineage (Nowack and Melkonian 2010), which only includes *Cardiosporidium* and *Nephromyces* (Muñoz-Gómez et al. 2019).

Endosymbiotic bacteria allow eukaryotes to exploit an enormous range of environments they would otherwise be unable to inhabit. Endosymbionts span a wide variety of taxa, from the *Buchnera* endosymbionts of aphids, which provide essential vitamins and amino acids, to the chemotrophic bacteria at the base of the deep-sea hydrothermal vent food chain. The diversity of prokaryotic metabolic pathways (McCutcheon, Boyd, and Dale 2019) drives the propensity of bacteria to colonize and exploit unusual habitats, including such extreme environments as radioactive waste (Fredrickson et al. 2004), highly acidic hot springs (Marciano-Cabral 1988), or even the inside of a host. In multicellular hosts, bacterial endosymbionts are frequently sequestered to specific structures or tissues, but in protists they must reside directly in the cytoplasm, making these associations far more intimate (Nowack and Melkonian 2010).

Though these interactions appear beneficial, endosymbiosis is rooted in conflict (Keeling and McCutcheon 2017; McCutcheon, Boyd, and Dale 2019). Many of the common endosymbiotic taxa, such as those within the order *Rickettsiales*, are closely related to pathogens. *Rickettsiales* is likely the sister taxon to the modern eukaryotic mitochondria (Fitzpatrick, Creevey, and McInerney 2006), and also contains *Wolbachia*, a genus of arthropod and nematode endosymbionts known to infect 25-70% of insects (Kozek and Rao 2007). Endosymbiosis and pathogenesis are closely related due to host cell invasion and persistence mechanisms (Keeling and McCutcheon 2017). However, the invading bacteria rarely see long term benefits from these interactions. Endosymbiont genomes are frequently found to be highly reduced due to the impact of Muller’s ratchet, in which population bottlenecks in vertically transmitted endosymbionts cause an accumulation of deleterious mutations over time (Moran 1996; McCutcheon and Moran 2012; Nowack and Melkonian 2010). With no gene flow between populations, endosymbionts are unable to recover from mutations and replication errors, which are more likely to occur in G/C rich regions, resulting in a characteristic A/T bias (McCutcheon, Boyd, and Dale 2019). The net impact of these forces is the creation of highly reduced, A/T rich genomes, which have convergently evolved in the majority of vertically transmitted endosymbiont lineages (Moran 1996; McCutcheon and Moran 2012; Keeling and McCutcheon 2017; McCutcheon, Boyd, and Dale 2019; Nowack and Melkonian 2010). Though the endosymbiont is fed and housed, it is also effectively incapacitated and permanently tied to its host.

Housing an endosymbiont is also costly for the host, and maintaining a foreign cell, rather than digesting or expelling it, indicates the endosymbiont confers a significant advantage. As part of a larger investigation of the Nephromycidae, here we focus on characterizing the role of the bacterial endosymbionts reported from *C. cionae* (Ciancio et al. 2008). Since *Cardiosporidium* and *Nephromyces* have maintained **α**-proteobacteria endosymbionts since before they diverged, we hypothesize this lineage of endosymbiont must provide metabolic functions of high value to its host apicomplexans. The maintenance of bacterial endosymbionts could be reducing host dependency and resource competition by providing novel biosynthetic pathways, thereby reducing virulence in this unique lineage.

## Materials and Methods

### Microscopy

Visual screens of *Ciona intestinalis* hemolymph were conducted using a 5% Giemsa/phosphate buffer stain with a thin smear slide preparation, as is commonly used to identify malarial infections (Moll et al. 2008). The filamentous life stage was identified during these screens due to its morphological similarity to *Nephromyces*. To confirm identity, three samples comprising 10-15 of the cell types of interest were manually picked and washed using stretched Pasteur pipettes and phosphate buffered saline. These samples were extracted, PCR amplified with *C. cionae* specific primers, and the resulting PCR product sequenced on the Sanger platform at the University of Rhode Island Genome Sequencing Center.

Fluorescence *in situ* hybridization (FISH) with 16S rRNA class specific probes was used to localize the bacterial endosymbionts as shown in Fig.1 (panels e and i). The hybridization was conducted as described in Paight et al., 2020.

**Figure 1:**
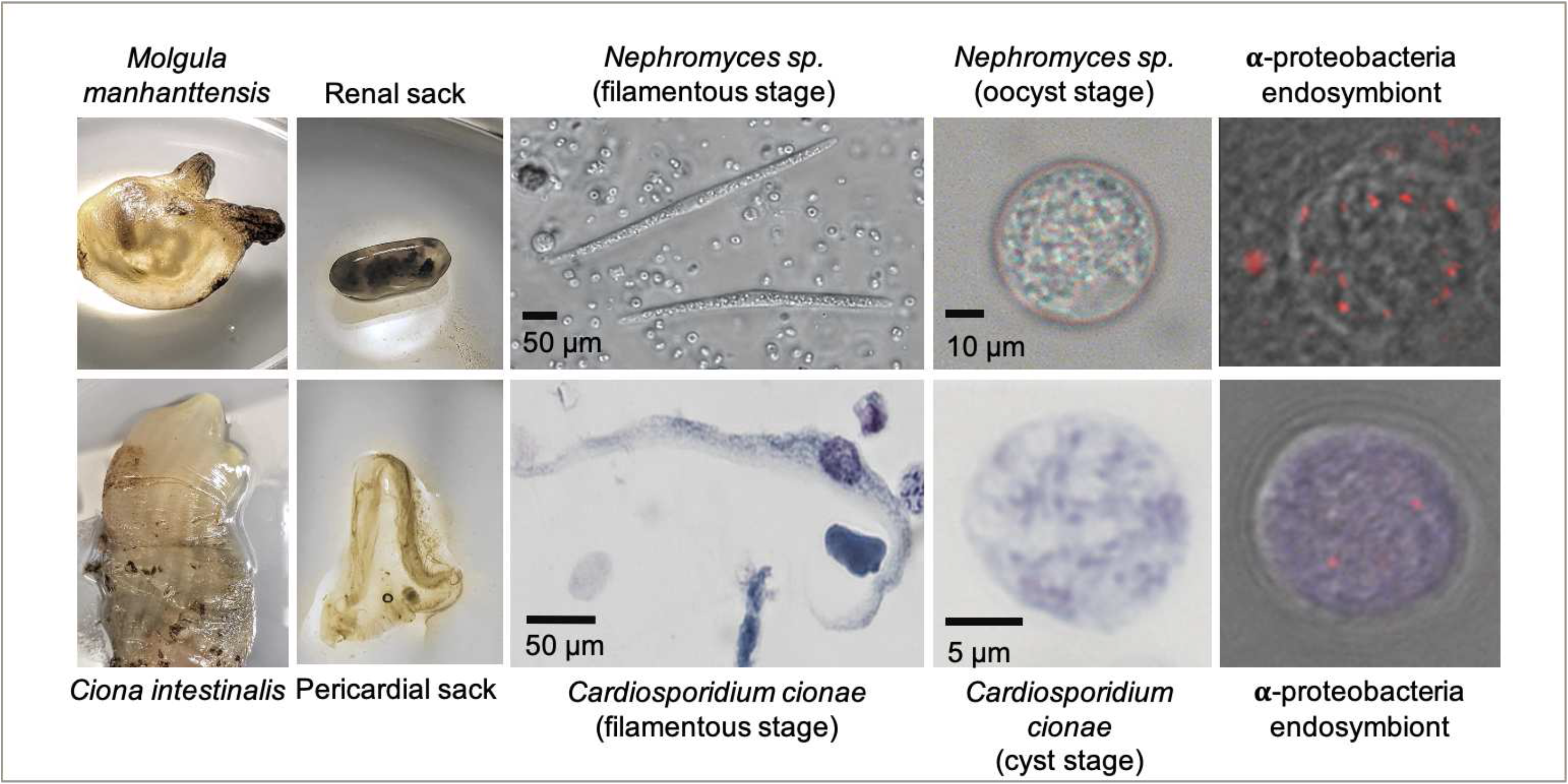
System overview of *Cardiosporidium cionae* and *Nephromyces* showing tunicate host (a and e), area of localization (b and f), filamentous life stage (c and g), oocyst life stage (d and h), and vertically transferred fluorescent *in situ* hybridization (FISH) labeled bacterial endosymbionts within the oocysts (e and i). Scale bars are approximations due to resizing of images. FISH was carried out according to the method in Paight et al. 2020.

### Extraction, Sequencing, Assembly, and Binning

The material for the *Cardiosporidium cionae* transcriptome was collected and isolated from wild *Ciona intestinalis* tunicates, as described in detail in Paight et al (2019). A sucrose density gradient was used to isolate *C. cionae* from tunicate hemolymph, and to enrich highly infected samples of hemolymph identified with microscopy. The gradient was composed of 20%, 25%, 30%, 35%, and 40% sucrose in phosphate buffer, loaded with approximately 5mL of hemolymph, and centrifuged in a swinging bucket rotor on 500 x g for 30 minutes, at 4°C (Paight et al. 2019). The 25% and 30% layers were then collected, pelleted, and washed with phosphate buffered saline, and stored at −80°C. RNA was extracted from the pellets and the highly infected samples used the Zymo Quick-RNA kit (Zymo Research LLC, Irvine, CA). Three samples with unfiltered hemolymph, hemolymph enriched with the 25% layer, and hemolymph enriched with the 30% layer were shipped on dry ice to the University of Maryland, Baltimore Institute for Genome Sciences, and multiplexed on a single lane of an Illumina HiSeq. These samples produced 92,250,706, 109,023,104, and 110,243,954 reads (Paight et al. 2019). They were assembled with Trinity/Trinotate v2.4.0 (Haas et al. 2014) and binned iteratively with OrthoFinder v2.3.3 (Emms and Kelly 2019) using a custom database of tunicates, Alveolates, and bacterial endosymbiont data to remove contamination from the host and environment.

For genomic sequencing, *C. intestinalis* were collected from Snug Harbor in South Kingstown, Rhode Island (41°23’13.4”N, 71°31’01.5”W) in August and September 2018, following the same protocol for dissection and needle extraction of the tunicate hemolymph from the pericardial sac. The sucrose density gradient described above was also used isolate *C. cionae* infected cells for genomic DNA, except that, in 3 of the 4 samples used, a 27% sucrose layer was substituted for the 25% layer to better capture *C. cionae* infected cells. In the fourth sample, a 30% layer was used. The layer of interest was centrifuged, collected, pelleted, and washed as described above, and in Paight et al. (2019). Filtered samples were used alone, rather than being incorporated into unfiltered hemolymph samples. Genomic DNA was immediately extracted using a 1% SDS lysis buffer, Proteinase K and phenol-chloroform extraction, followed by an overnight ethanol precipitation at −20 °C. Samples were assessed for quality and concentration with gel electrophoresis, nanodrop, and Qubit (broad range), and then stored at −20 °C.

Samples from four separate gradient columns were individually prepared at the University of Rhode Island Genome Sequencing Center, and the resulting libraries run on a single lane of an Illumina HiSeq4000 at the University of Maryland, Baltimore Institute for Genome Sciences. These libraries were independently trimmed and assessed for quality using Trimmomatic v0.36 and FastQC v0.11.8 before being pooled and assembled with SPAdes v3.13.0 on the Brown University OSCAR server (Bolger, Lohse, and Usadel 2014; “FastQC A Quality Control Tool for High Throughput Sequence Data” n.d.; Bankevich et al. 2012).

The SPAdes metagenomic assembly was binned by assigning taxonomy to contigs with CAT (von Meijenfeldt et al. 2019). *Rickettsiales* sequences were confirmed using MetaBAT (Kang et al. 2015), and the resulting contigs inspected for contamination and reassembled with Geneious v9.1.8 (“Geneious | Bioinformatics Software for Sequence Data Analysis” 2017). Additional apicomplexan sequences were identified by mapping trimmed and binned transcriptomic reads to the full metagenomic assembly using Bowtie2 v2.3.5.1, and contig coverage calculated with the bedtools v2.26.0 genomecov function (Langmead and Salzberg 2012; Quinlan and Hall 2010). The resulting file was sorted with R to extract contigs with greater than 50% coverage of *C. cionae* transcripts. Both the *C. cionae* and **α**-proteobacterial endosymbiont genomic assemblies were trimmed to a minimum length of 1kb, as contigs smaller than this were unlikely to be reliably binned. Genome assembly graphs were also visualized with Bandage v0.8.1 (Wick et al. 2015) and clusters of interest were identified with BLAST. The **α**-proteobacteria cluster was identified with BLAST, exported, and compared to the CAT binned bacterial assembly with average nucleotide identity estimations (Rodriguez and Konstantinidis 2016).

Organellar assemblies for both the apicoplast and the mitochondrion were generated with NOVOPlasty v3.7.2 (Dierckxsens, Mardulyn, and Smits 2016). The seed sequences for these assemblies were located using the apicoplast genomes of *Nephromyces* (Muñoz-Gómez et al. 2019), and Sanger sequences of the *C. cionae* cytochrome C oxidase subunit one (COX-1) gene generated with PCR with local BLASTN databases (Madden 2003).

### Gene Prediction and Annotation

Annotation of the **α**- proteobacteria endosymbiont genome and *C. cionae* mitochondria was carried out using Prokka v1.14.5 (Seemann 2014). Closely related *Rickettsia* genomes (as indicated by the preliminary 16S phylogeny) were retrieved from NCBI and used to generate a custom database for **α**- proteobacteria genome annotation (Seemann 2014) (Table S1). Annotation of the apicoplast was carried out in Geneious v9.1.8 using a custom database of *Nephromyces* apicoplast annotations (Muñoz-Gómez et al. 2019). Inverted repeat regions were identified with Repeat Finder plugin.

*Cardiosporidium cionae* genomic contigs were annotated with the MAKER v2.31.10 pipeline (Holt and Yandell 2011). Repeats were soft masked using RepeatMasker v4.0.9 (Smit, Hubley, and Green 2013). *Ab initio* predictions and species training parameters were generated with both WebAugustus (Hoff and Stanke 2013) and SNAP-generated hidden markov models (Korf 2004). This process was repeated iteratively, and the AED values indicating the fit of gene prediction to the model were analyzed to ensure high quality predictions. Predicted proteins from both organisms were functionally classified with the Kyoto Encyclopedia of Genes and Genomes (KEGG) and NCBI BLASTP v2.7.0+ (Kanehisa et al. 2017; Madden 2003). Coding sequences were searched for homologous domains with InterProScan (Mitchell et al. 2019). Individual genes of interest were screened using BLAST databases.

### Analysis

Completeness of the *C. cionae* genome and transcriptome were assessed with BUSCO using the eukaryotic database (Simão et al. 2015). Homologs identified as multicopy by BUSCO were manually screened with NCBI-BLAST to confirm they did not represent contamination in the finished assembly. Proteins were annotated using orthologues from EuPathDB, PFAM, Kegg, and Interpro. Transcripts with multiple predicted isoforms in the transcriptome were filtered and selected based on completeness, Interpro score, and length. Completeness and contamination of the *C. cionae* **α**- proteobacteria genome was assessed with the Microbial Genome Atlas (MiGA) and CheckM (Parks et al. 2015; Rodriguez-R et al. 2018). Candidate pseudogenes were located with Pseudofinder with default parameters, using a custom BLAST database of proteins from the 785 complete alphaproteobacterial genomes available on NCBI (Syberg-Olsen et al. 2018). Visual representations of the metabolic pathways were constructed for both *C. cionae* and the **α**- proteobacteria using functional annotations from KEGG (Kanehisa et al. 2017). Visual representations of the **α**- proteobacteria were generated using Circos (Krzywinski et al. 2009) with annotation data from Prokka and functional annotations from KEGG.

In addition to functional comparisons using KEGG annotations, the **α**- proteobacterial endosymbiont genome was also compared to the *Nephromyces* endosymbiont with similarity estimations and orthologous gene content. Similarity was compared with average nucleotide identity and average amino acid identity calculations using the web-based ANI and AAI calculator (Rodriguez and Konstantinidis 2016).

Orthologous gene content comparisons between the *C. cionae* **α**-proteobacteria and all of the endosymbionts in the *Nephromyces* system was carried out with OrthoFinder v2.3.3 (Emms and Kelly 2019). The resulting overlaps were calculated using the R package limma (Ritchie et al. 2015), and the final figure generated with Venn Diagram (Chen and Boutros 2011), also in R. Functional gene overlap was based on KEGG annotations and generated using the same R packages.

### Phylogenetics

Bacterial phylogenies were constructed using the predicted taxonomy from MiGA, which assigned the endosymbiont to the class alphaproteobacteria with a p value of 0.25 (Parks et al. 2015; Rodriguez-R et al. 2018). To confirm this result, all complete bacterial proteome accessions belonging to this class were retrieved from the NCBI database (712 in total). These data were searched using the **α**-proteobacteria HMM single copy gene set comprised of 117 proteins, aligned, and the tree constructed using the GToTree workflow (Lee 2019; “Accelerated Profile HMM Searches” n.d.; Hyatt et al. 2010; Price, Dehal, and Arkin 2010; Edgar 2004; “TaxonKit - NCBI Taxonomy Toolkit” n.d.; Capella-Gutiérrez, Silla-Martínez, and Gabaldón 2009).

The apicoplast encoded genes of *C. cionae* were added to the dataset used in (Muñoz-Gómez et al. 2019) to confirm monophyly with *Nephromyces*, previously indicated with COI and 18S gene trees. Protein homologs were identified using local BLAST-P searches and concatenated with the existing dataset. These sequences were aligned with MAFFT v7, trimmed in Geneious, and concatenated (Madden 2003; Katoh and Standley 2013). Species phylogeny was inferred with Maximum Likelihood using IQ-TREE (v1.6) and the LG+G model. Statistical support at branches was estimated using ultrafast bootstrap (1000) and aLRT (1000) (Nguyen et al. 2015).

### Parameters

The specific scripts and settings used for bioinformatic analysis of *C. cionae* and its endosymbiont have been deposited in a publicly accessible GitHub repository (github.com/liz-hunter/cardio_project).

## Results

### Cardiosporidium cionae

Genomic sequencing of the pooled *C. cionae* libraries yielded a total of 320,000,000 paired reads. After trimming and assembly, this resulted in 656,251 contigs, 176,701 of which were larger than 1kb. Binning with CAT resulted in 3,641 contigs assigned to the superphylum Alveolata. Contigs assigned to Dinophyceae and Ciliophora were removed, leaving 2754 contigs, and 1790 of these contigs were larger than 1kb. The RNA-seq assisted coverage-based binning added an additional 935 contigs, 423 of which were unique and larger than 1kb. Further manual curation using OrthoFinder eliminated 7 additional contigs. This resulted in a total of 2,206 contigs assigned to *C. cionae.* Of the remaining 174,496 contigs, 221 were assigned to the order Rickettsiales, and 147,793 contigs assigned to the class Ascidiacea (tunicate). The *C. cionae* genome assembly is 57Mb in total, with an N50 of 54.04kb, and a G/C content of 34.4% (Table 1). This is smaller than some apicomplexan genomes such as coccidian *Toxoplasma gondi* (80Mb), but considerably larger than haemosporidian *Plasmodium falciparum* (22.9Mb) and the highly reduced *Cryptosporidium parvum* (9Mb) (Sibley and Boothroyd 1992; Abrahamsen 2004). Gene prediction resulted in 4,674 proteins (Table 1). The binned transcriptome assembly yielded a total of 15,077 proteins assigned to *C. cionae*, including all isoforms. When filtered to remove redundancy, this dataset was reduced to 6,733 unique proteins.

**Table 1:** Statistics for the genomic and transcriptomic datasets presented. MiGA was used to assess the bacterial genome completeness, and BUSCO was used to assess the *Cardiosporidium cionae* genome and transcriptome.

| Assembly | Contigs | Size<br>(Mb) | N50<br>(kb) | N90<br>(kb) | G/C<br>(%) | Proteins | Completeness<br>(%) | Duplication<br>(%) |
| --- | --- | --- | --- | --- | --- | --- | --- | --- |
| *C. cionae* (DNA) | 2,213 | 57 | 54.12 | 15.21 | 34.4 | 4,692 | 63.7 | 13.9 |
| α-proteobacteria | 29 | 1.05 | 250.39 | 36.72 | 29.1 | 906 | 91.9 | 1.9 |
| *C. cionae* (RNA) | - | - | - | - | - | 6,733 | 60.0 | 4.6 |

The final binned *C. cionae* genome assembly is estimated to be 63.7% complete by BUSCO, with 13.9% duplication. The transcriptome is slightly more complete, with a BUSCO estimate of 68.3% complete orthologs, and 12.5% partial (Paight et al. 2019). When the isoforms were filtered for annotation, this completeness value dropped slightly to 60.0% with 4.6% duplication (Table 1). Despite this, the *C. cionae* assembly contains genes from all of the expected core biosynthetic pathways for a haematozoan. *Cardiosporidium* has a suite of basic metabolic pathways including complete or nearly complete functional predictions for glycolysis, gluconeogenesis, pyruvate oxidation, the pentose phosphate cycle, and the citric acid cycle (Fig. 2, Table S2). It also encodes a handful of unexpected pathways, including the entire *de novo* IMP biosynthetic pathway. *Cardiosporidium* contains the genes for fatty acid biosynthesis and elongation in the endoplasmic reticulum, as well degradation to produce acetyl-CoA (Fig. 2, Table S2).

**Figure 2:**
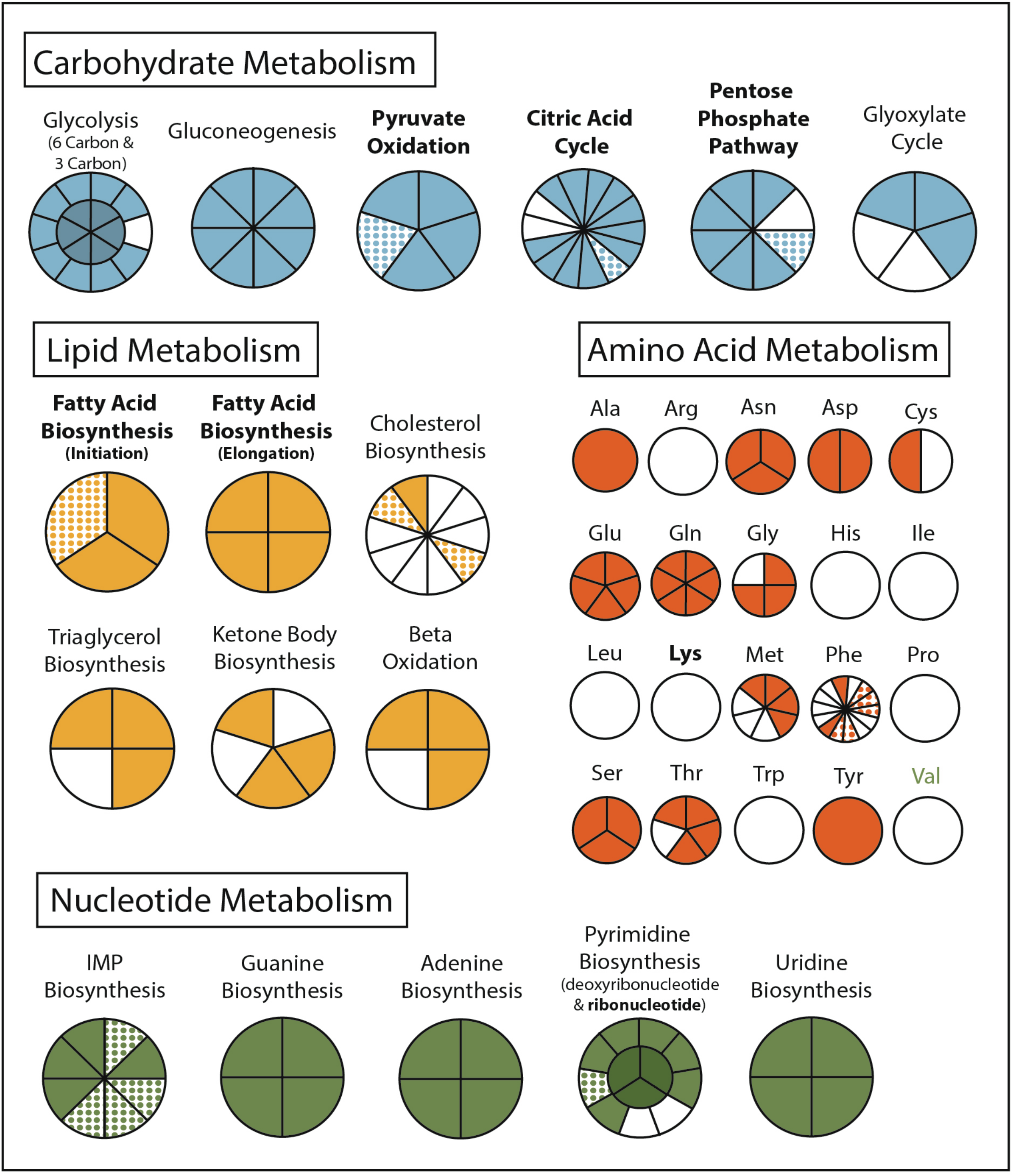
Overview of the metabolism of *Cardiosporidium cionae*. Solid colors indicate genomic protein homologs, dots show where homologs were only found in the transcriptome data, and bold pathways represent bacterial endosymbiont contributions. This figure corresponds to the genome and transcriptome information in supplementary table 2.

*Cardiosporidium* also encodes the majority of the pathway of triacylglycerol biosynthesis, and partial pathways for cholesterol and ketone body synthesis. It completely lacks any evidence of biosynthetic genes for eight of the twenty-one amino acids but does encode amino acid conversion pathways that other Hematozoa lack. These include the conversion of phenylalanine to tyrosine, and homocysteine to methionine (Fig. 2, Table S2). Additionally, *C. cionae* is able to generate serine from multiple sources (glycerate-3P and glycine), as well as degrade it to pyruvate. The genomic data we recovered only encodes partial pathways for riboflavin, and heme synthesis, and also lacks genes for biotin, thiamine, ubiquinone, and cobalamin synthesis. However, we identified both C5 and C10-20 isoprenoid biosynthesis pathways. This genome also supports the presence of the purine degradation pathway previously identified in the transcriptome of *Nephromyces* (Paight et al. 2019).

Visual screens with thin-smear Giemsa staining indicate that *C. cionae* maintains a very low density inside its host. These microscopy screens further revealed the presence of a large, extracellular filamentous life stage analogous to the filamentous life-stage in *Nephromyces* (Fig. 1). Single cell isolation, extraction, and PCR confirmed these cell types were indeed a life-stage of *C. cionae*.

Phylogenetic analysis of the apicoplast encoded proteins supported the monophyly of *Nephromyces* and *C. cionae* (Fig. 3). This analysis differs with the placement results for *Nephromyces* published by Muñoz-Gómez et al. 2019, due to maximized data and the omission of early branching taxa. This taxon sampling caused Nephromycidae to branch outside of the Hematozoa. The *C. cionae* apicoplast is structurally very similar to those of *Nephromyces* in terms of gene content, size, and organization (Fig. S1).

**Figure 3:**
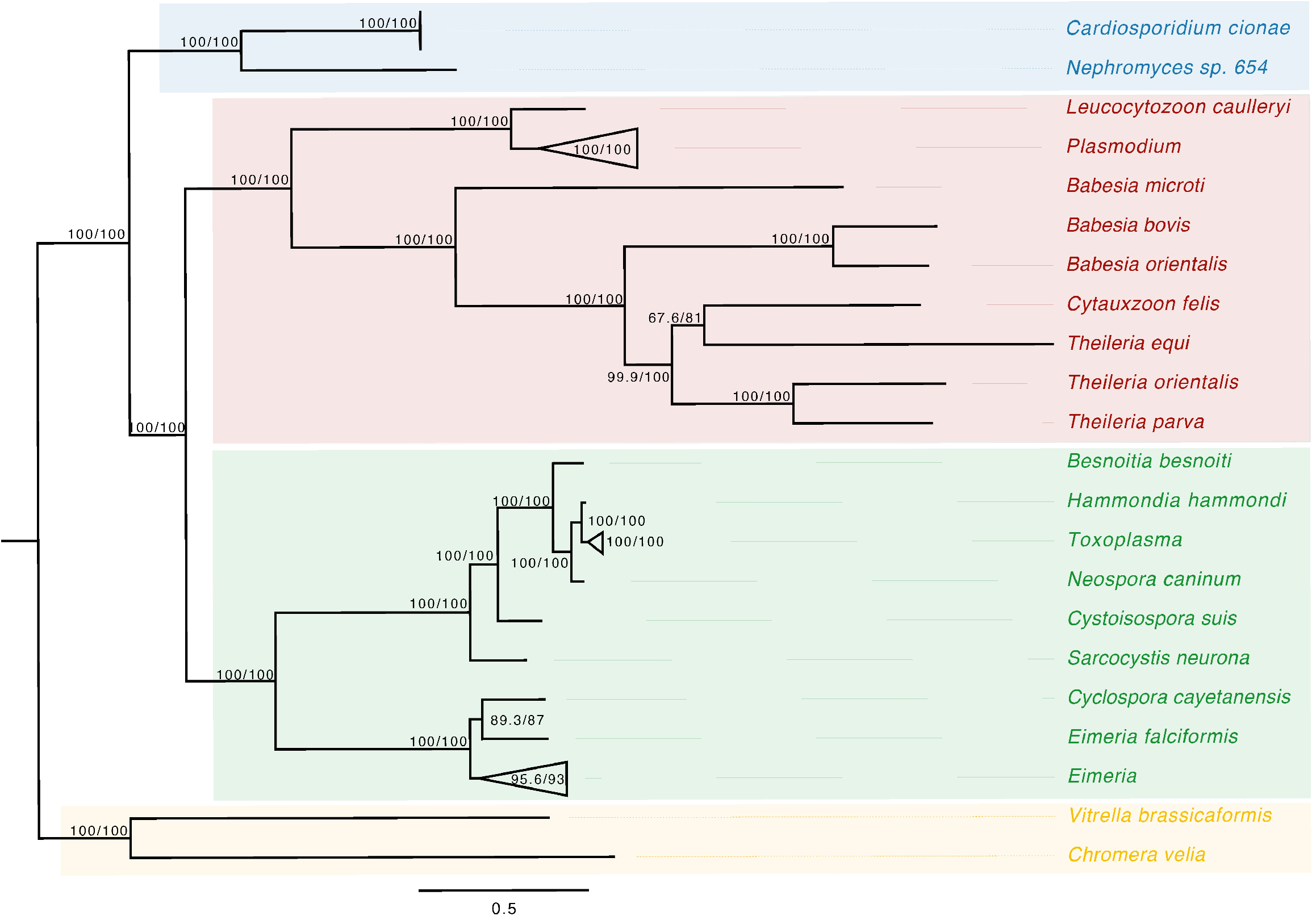
Apicoplast phylogeny created using a modified dataset provided by Muñoz-Gómez et al. 2019, showing the monophyly of *C. cionae* and *Nephromyces*. The complete, circularized *C. cionae* apicoplast recovered from the genomic dataset, and *Nephromyces* apicoplast sp. 654, are shown in blue. Statistical support was estimated using aLRT (1000) and ultrafast bootstrap (1000) (Nguyen et al. 2015) and these values are shown in this order on the nodes.

### **α**- proteobacteria

Of the contigs assigned to Rickettsiales, 45 were larger than 1kb. Reassembly yielded 31 contigs, and manual curation resulted in a final 29 contigs. The final **α**-endosymbiont assembly is 1.05Mb in total, with an N50 of 250.39kb, and a G/C content of 29.1%. Gene prediction and annotation resulted in 906 proteins (Table 1). The Bandage cluster was shown to have an average nucleotide identity (two-way ANI) of 99.95% (SD.81%) based on 4,878 fragments when compared with the CAT binned assembly. This provided independent validation for the bacterial genome assembly binning. The **α**-endosymbiont assembly is estimated to be 91.9% complete with 1.8% contamination by MiGA, and 95.5% complete with 2.1% contamination by CheckM.

Characteristic of bacterial endosymbionts, it has a low G/C content and high coding density (Fig. 4). This organism is predicted to encode just 906 genes by Prokka (Table 1), and 37 of which were identified as candidate pseudogenes by Pseudofinder. These genes were largely hypothetical proteins, but 15 genes were also identified as likely being nonfunctional. These included a permease, transposase, thioesterase, phosphodiesterase, and multiple transferases. Pseudofinder also joined 13 ORFs, leaving only 865 predicted functional genes. With so few functional genes, it is not surprising that this **α**-proteobacteria has a sparse number of complete metabolic pathways (Fig. 4, Fig. 2). The genome is slightly smaller than closely related alphaproteobacterial endosymbionts, such as *Candidatus* Phycorickettsia trachydisci sp. nov. (1.4MB), *Orientia tsutsugamushi* (2MB), and other protist associated Rickettsiales lineages (1.4-1.7MB) (Yurchenko et al. 2018; Nakayamak et al. 2010; Muñoz-Gómez et al. 2019). The **α**- proteobacteria genomes in both *C. cionae* and *Nephromyces* encode pathways for the biosynthesis of fatty acids, pyrimidines, lipoic acid, heme, glutamine, lysine, ubiquinone, and the citric acid cycle. Only the *C. cionae* **α**- proteobacteria maintains the genes for asparagine biosynthesis, glycolysis, and the pentose phosphate pathway (Fig. S2), while only the *Nephromyces* **α**- proteobacteria can complete glutamic acid biosynthesis.

**Figure 4:**
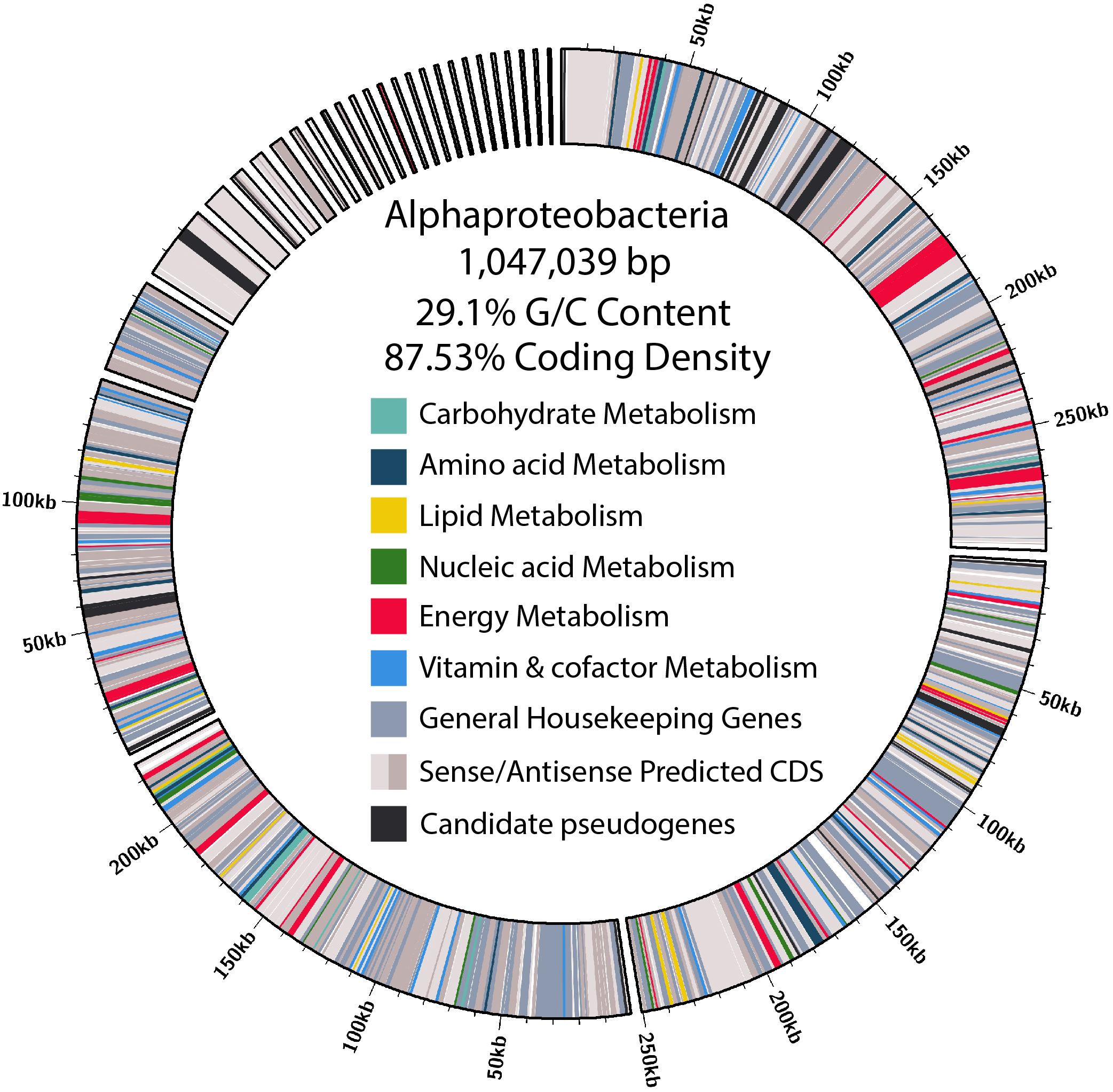
Overview of size, contig distribution, coding density, and annotations of major functional categories of genes in the **α**-proteobacteria endosymbiont genome.

When the **α**- proteobacteria in both *Cardiosporidium* and *Nephromyces* were compared for similarity, the results showed these taxa were too divergent to be compared with average nucleotide identity (ANI), and they were instead compared with average amino acid identity (AAI). A two-way AAI analysis of 656 proteins showed 47.61% (SD:12.51%) similarity between these genomes, which is consistent with the phylogenetic analysis that indicates considerable evolutionary distance between these two taxa. This multigene phylogeny of the **α**-endosymbionts is congruent with the preliminary 16S gene trees, which places these species in the order Rickettsiales. They belong to the family Rickettsiaceae and are sister to the genus *Rickettsia* (Fig. 5).

**Figure 5:**
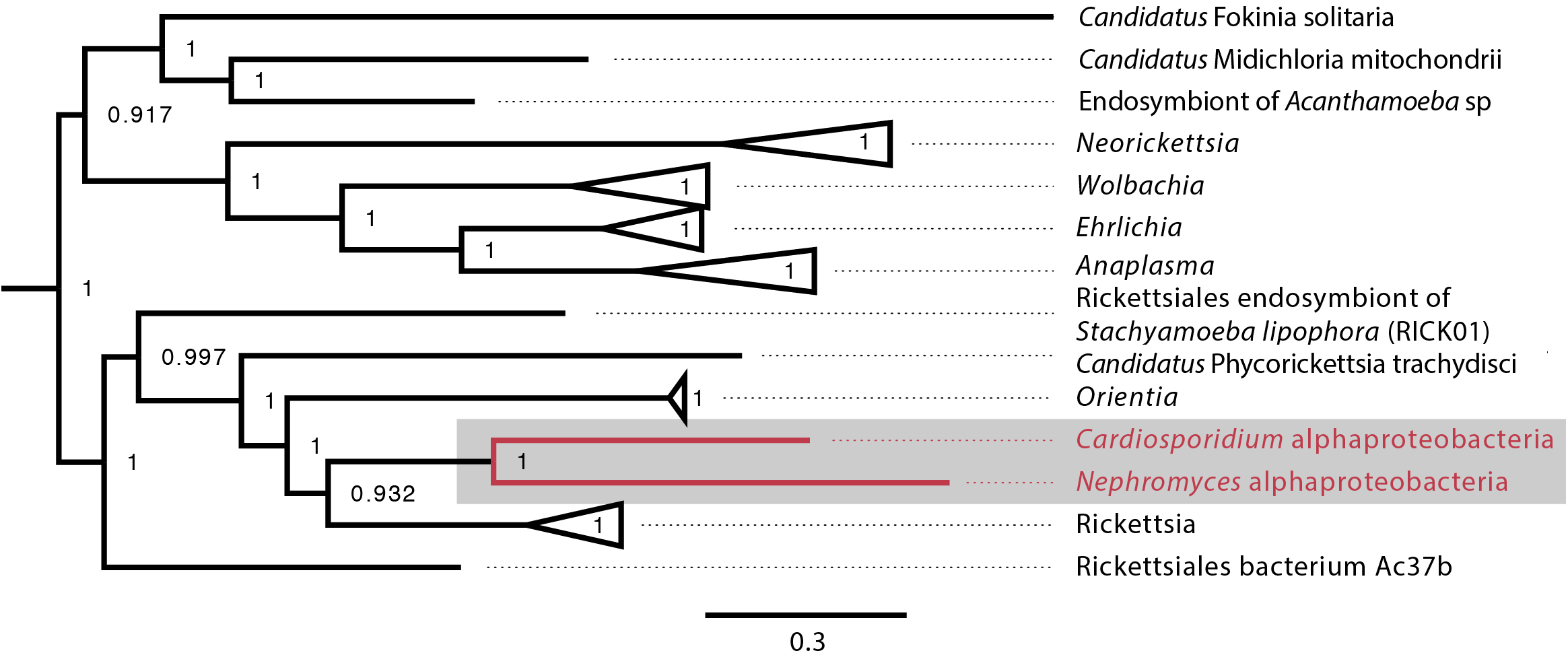
Alphaproteobacteria phylogeny created with GToTree pipeline (117 concatenated genes) including all sequenced alphaproteobacteria published on NCBI and both the *C. cionae* and *Nephromyces* **α**-endosymbionts (in red). Bootstrap support is shown as a decimal value on the nodes.

Ortholog comparisons between the **α**-endosymbionts indicate these taxa share the majority of their core functions, but the *Cardiosporidium* system **α**-endosymbiont maintains more unique genes. This taxon also shares greater ortholog and functional overlap with the two additional endosymbionts present in the *Nephromyces* system: betaproteobacteria and Bacteroides (Fig. S2).

### Data Availability

All data associated with this project is deposited in GenBank under the BioProject PRJNA664590. The *Cardiosporidium cionae* whole genome shotgun projected has been deposited under the accession JADAQX000000000, and the alphaproteobacterial endosymbiont genome is deposited under the accession JADAQY000000000, and the transcriptome is deposited under the accession GIVE00000000. The versions described in this paper are versions JADAQX010000000, JADAQY010000000, and GIVE01000000.

## Discussion

Metabolically, *C. cionae* is similar to other sequenced hematozoans. However, it also encodes some unusual pathways. *Cardiosporidium cionae*, like *Nephromyces*, encodes the *de novo* purine biosynthesis pathway (Fig. 2, Table S2), which has been lost in all other sequenced apicomplexans (Janouskovec and Keeling 2016). These genes resolve with *Nephromyces, Vitrella brassicaformis* and dinoflagellates such as *Crypthecodinium cohnii* in phylogenetic analysis, demonstrating this was not a recent horizontal gene transfer event (Paight et al., 2020). Instead, these data indicate that both genera within Nephromycidae have maintained the ancestral pathway found in free-living Chromerids, and the genes for purine biosynthesis have been lost independently in all other apicomplexan lineages. *De novo* biosynthesis of purines in *C. cionae* and *Nephromyces* reduces dependence on preformed purine metabolites from their respective hosts, potentially enabling the persistence of the extracellular life stages in both of these lineages. *Nephromyces* and *C. cionae* are also able to degrade purines (Paight et al. 2019), and we suspect this aspect of their metabolism is related to the physiology of tunicates, which are incapable of metabolizing uric acid, a purine waste product. Even though the tunicate hosts are unable to degrade uric acid, they inexplicably accumulate it (Nolfi 1970; Lambert et al. 1998).

Whereas the complete pathways for pentose phosphate cycle, citric acid cycle, and gluconeogenesis mirror other hematozoans (Fig. 2, Table S2), *C. cionae* also encodes a handful of genes that suggest it is able to produce glyoxylate. Paight et al. (2019) reported transcripts for a number of peroxisomal proteins in both *C. cionae* and *Nephromyces*, and predicted a novel metabolic pathway. Despite their numerous metabolic similarities, *C. cionae* and *Nephromyces* appear to have distinct pathways for central carbon metabolism (specifically the citric acid cycle), and part of the closely linked glyoxylate cycle. Both *Nephromyces* and *C. cionae* possess a uniformly highly expressed purine degradation cycle that converts ureidoglycolate to glyoxylate using a novel amidohydrolase, and generate glycine and serine. However, only *Nephromyces* can feed glyoxylate back into the citric acid cycle using malate synthase (Paight et al. 2019). This pathway is a product of the unusual renal sac environment where *Nephromyces* makes its home, which contains an abundance of uric acid sequestered by the host tunicate. In *C. cionae*, malate synthase is conspicuously absent in both the genome and transcriptome, indicating the carbon cycling in these closely related organisms is likely distinct, and potentially one of the differences that accounts for the virulence disparity between *C. cionae* and *Nephromyces.* However, the list of differences, which also includes host species and organellar localization, is relatively short. Though it was known that these taxa have similar life history traits (Ciancio et al. 2008; Saffo et al. 2010), these genomic data suggest their morphology and metabolism are also remarkably similar.

*Cardiosporidium cionae* and *Nephromyces* (Nephromycidae) branch within Haematozoa, a group of obligate, parasitic, intracellular apicomplexans (Muñoz-Gómez et al. 2019; Mathur et al. 2019). All previously described members of Haemotozoa, and sister taxon Coccidia, are intracellular and obligately parasitic. Despite their phylogenetic position within an obligately intracellular clade, members of the Nephromycidae have large, filamentous, extracellular life stages (Fig. 1). *Nephromyces* is completely extracellular (Saffo and Nelson 1982), while *C. cionae* has both intracellular and extracellular life stages. Though morphologically similar to the more basal gregarine apicomplexans (Rueckert et al. 2015), these groups are phylogenetically distant. The Nephromycidae have evolved from intracellular ancestors, and has transitioned to the extracellular environment. In *Nephromyces*, this transition is complete, whereas *C. cionae* has both intracellular and extracellular life stages. We believe that extracellularity in this group is related to another unusual characteristic: the maintenance of bacterial endosymbionts in both *C. cionae* and *Nephromyces*.

The maintenance of monophyletic **α**-endosymbionts in both the *C. cionae* and *Nephromyces* lineages indicates that this endosymbiont is providing something vital to the system. However, at first glance, these endosymbionts are contributing very little to their host apicomplexans. Like its counterpart in *Nephromyces*, the *C. cionae* **α**-endosymbiont contains only a handful of biosynthetic pathways (Fig. 4). Overall, the **α**-endosymbiont in *C. cionae* does contain more unique orthologs and functional genes when compared with its counterpart in the *Nephromyces* system (Fig. S2). Primarily, these unique genes are related to energy metabolism (Fig. 4, Fig. S2), and their presence is likely a result of the heightened evolutionary pressure to maintain critical genes in a system with a single endosymbiont, compared to the three types of endosymbionts present in *Nephromyces* communities. The **α**-endosymbiont encoded pathways for energy and carbon cycling, while possibly advantageous to *C. cionae*, are likely not critical contributions because they can be completed by the apicomplexan, independent of the endosymbiont. The maintenance of an endosymbiont is costly, and it is unlikely to be preserved for a redundant function (McCutcheon and Moran, 2012).

A handful of pathways have been maintained in both **α**-endosymbiont lineages, and are also absent in the host apicomplexans. The only apparently critical functions that cannot be replaced by the apicomplexan metabolism are lysine biosynthesis, and lipoic acid biosynthesis. Lysine is an essential amino acid, and plays an important role in protein biosynthesis. Lysine is an essential media component for the growth of *P. falciparum* and is predicted to be scavenged from the host by *T. gondii* (Tymoshenko et al. 2015; Schuster 2002). Lysine biosynthesis is also absent in the *Nephromyces* genome and transcriptome (Paight et al., 2020). Like *Nephromyces, T. gondii* and *P. falciparum*, our data indicate *C*. *cionae* cannot synthesize its own lysine and is dependent on host scavenging. Though we cannot exclude the possibility that *C. cionae* encodes lysine biosynthesis with an incomplete genome, based on the genomes of other apicomplexans, lysine is likely absent within Nephromcyidae. Lysine is also essential for the host tunicate, *Ciona intestinalis* (Kanehisa et al. 2017), and both organisms requiring environmental sources of lysine puts them in constant competition for the resource. *Cardiosporidium cionae* appears to have circumvented this conflict by maintaining a bacterial endosymbiont that contains the pathway for *de novo* lysine biosynthesis. Rather than compete with the host tunicate for lysine, *C. cionae* cultivates an intracellular source for the essential amino acid, reducing host dependency and potentially virulence.

Lipoic acid is an aromatic sulfur compound that is an essential cofactor for a series of vital metabolic functions. These include the citric acid cycle and alpha keto dehydrogenase complexes, such as the pyruvate dehydrogenase complex and the glycine conversion system. In eukaryotes, lipoic acid is exclusively localized to the mitochondria and the plastid. Apicomplexans localize lipoic acid biosynthesis to the apicoplast, having lost the mitochondrial pathway after the acquisition of the plastid (Crawford et al. 2006). Instead, an alternative scavenging pathway is used to produce the lipoic acid required for the citric acid cycle and glycine conversion system in the mitochondria, and both the scavenging and biosynthetic pathways are considered essential (Günther et al. 2005). Functional studies have shown that when the lipoic acid biosynthetic pathways are knocked out, *P. falciparum* will compensate by scavenging more lipoic acid from the host and shuttling it to the apicoplast (Günther et al. 2007). Similarly, *T. gondii* growth is inhibited by lipoate-deficient media, suggesting scavenging is essential (Crawford et al. 2006). Metabolic modeling also indicates that even apicomplexans that maintain this pathway require supplemental lipoic acid from their host organisms (Blume and Seeber 2018). Though lipoic acid is produced by the host tunicates, we speculate that there is limited availability for an extracellular organism because it is both produced and used in the mitochondria. This likely means *C. cionae* is dependent on this **α**-endosymbiont for the production of key compounds such as lipoic acid, for the persistence of a stable extracellular life stage. In this way, maintaining the **α**-endosymbiont as an internal cofactor source further reduces resource competition between *C. cionae* and its host.

The Nephromycidae have evolved from a clade of an obligately parasitic intracellular apicomplexans, and have transitioned to a mostly extracellular lifestyle. We hypothesize that, by obtaining bacterial endosymbionts, these apicomplexans have acquired metabolic capabilities that enabled this transition. Though *Nephromyces* shares an **α**-endosymbiont lineage with *C. cionae* (Fig. 5), it also has betaproteobacteria and Bacteroides endosymbionts. With this bacterial taxonomic diversity comes metabolic diversity, and though the **α**-endosymbiont in *C. cionae* has more unique functional proteins and orthologs than its counterpart, this is dwarfed by the number of unique proteins and orthologs contributed by the two additional taxa present in the *Nephromyces* system (Fig. S2). We believe the sole endosymbiont in *C. cionae* provides a dedicated source of the essential metabolites lysine and lipoic acid, which likely reduces competition with its host compared to its haematozoan relatives, and makes extracellular life stages possible. In this way, *Cardiosporidium cionae* represents a potential intermediate in the transition to mutualism, that has been described in *Nephromyces* (Saffo et al. 2010).

---

**Supplementary Table 1:** Closely related reference genomes used for annotation of the *Cardiosporidium cionae* **α**-endosymbiont, downloaded from GenBank, refseq.

| ***Rickettsia* reference genomes** | **RefSeq Accession** |
| --- | --- |
| Rickettsia conorii str. Malish 7 | GCF_000007025.1 |
| Rickettsia felis str. Pedreira | GCF_000964665.1 |
| Rickettsia japonica YH | GCF_000283595.1 |
| Rickettsia massiliae MTU5 | GCF_000016625.1 |
| Rickettsia peacockii str. Rustic | GCF_000021525.1 |
| Rickettsia prowazekii str. Breinl | GCF_000367405.1 |
| Rickettsia typhi str. Wilmington | GCF_000008045.1 |
| Rickettsia bellii RML369-C | GCF_000012385.1 |

**Supplementary Table 2:**
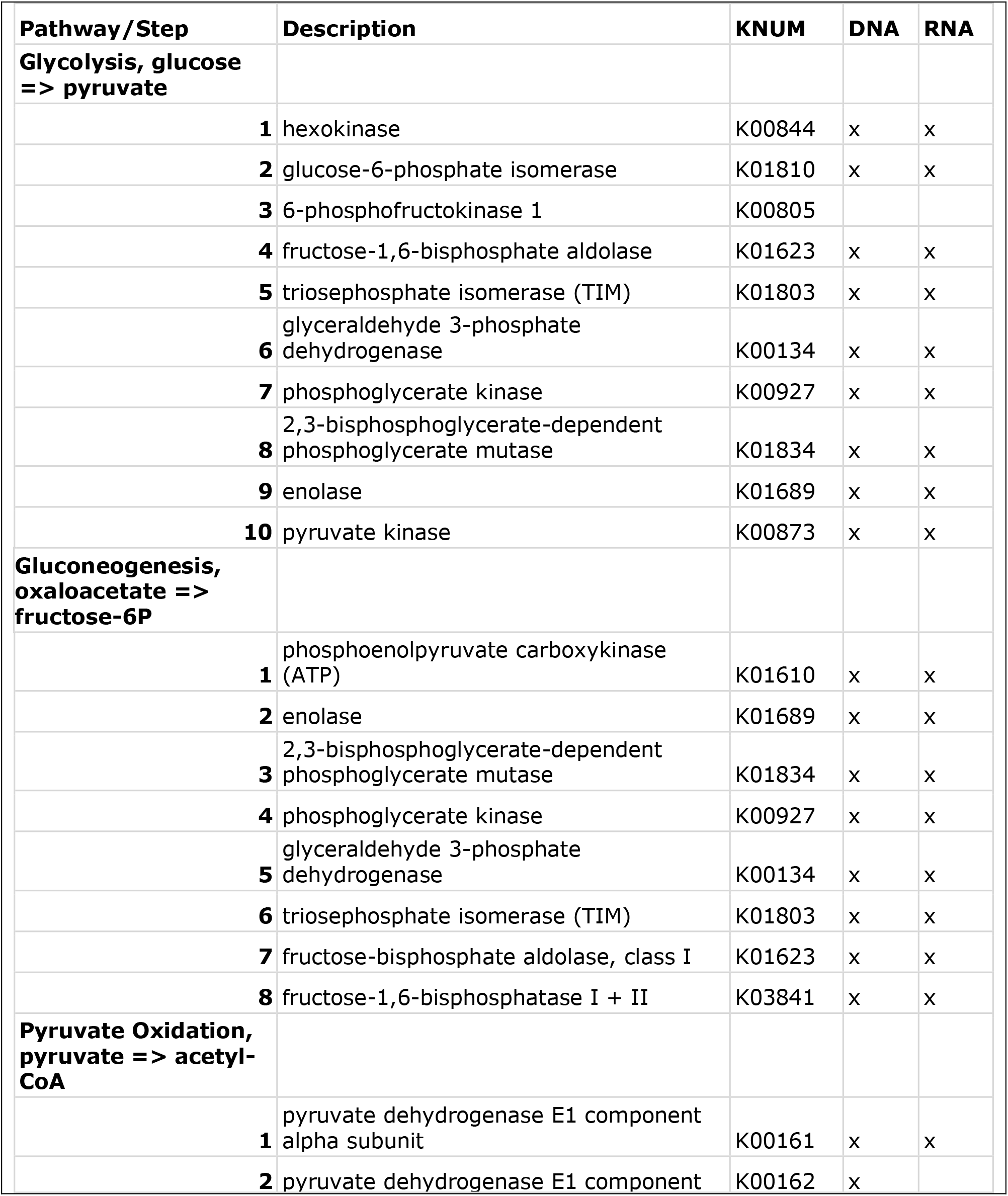
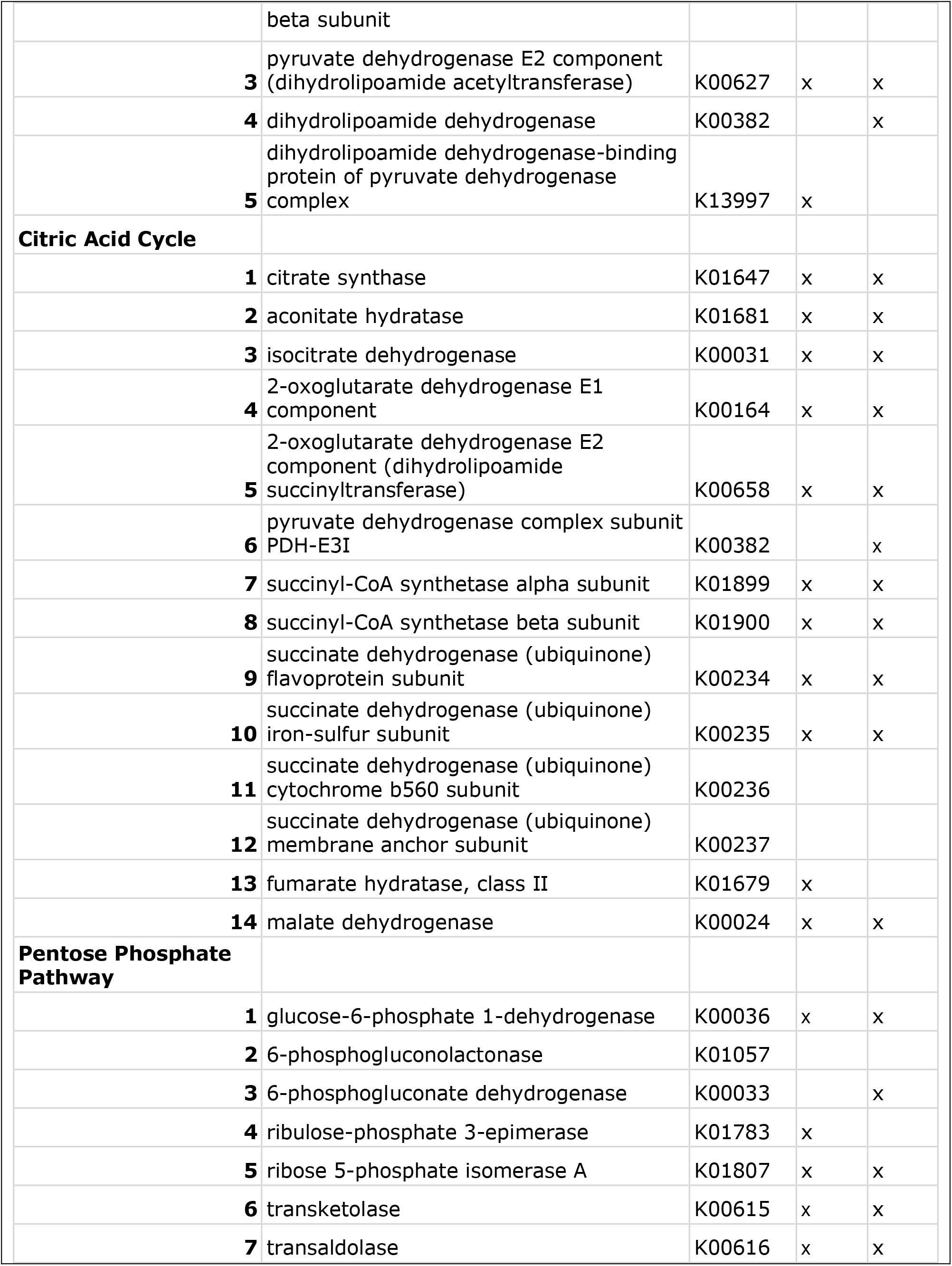
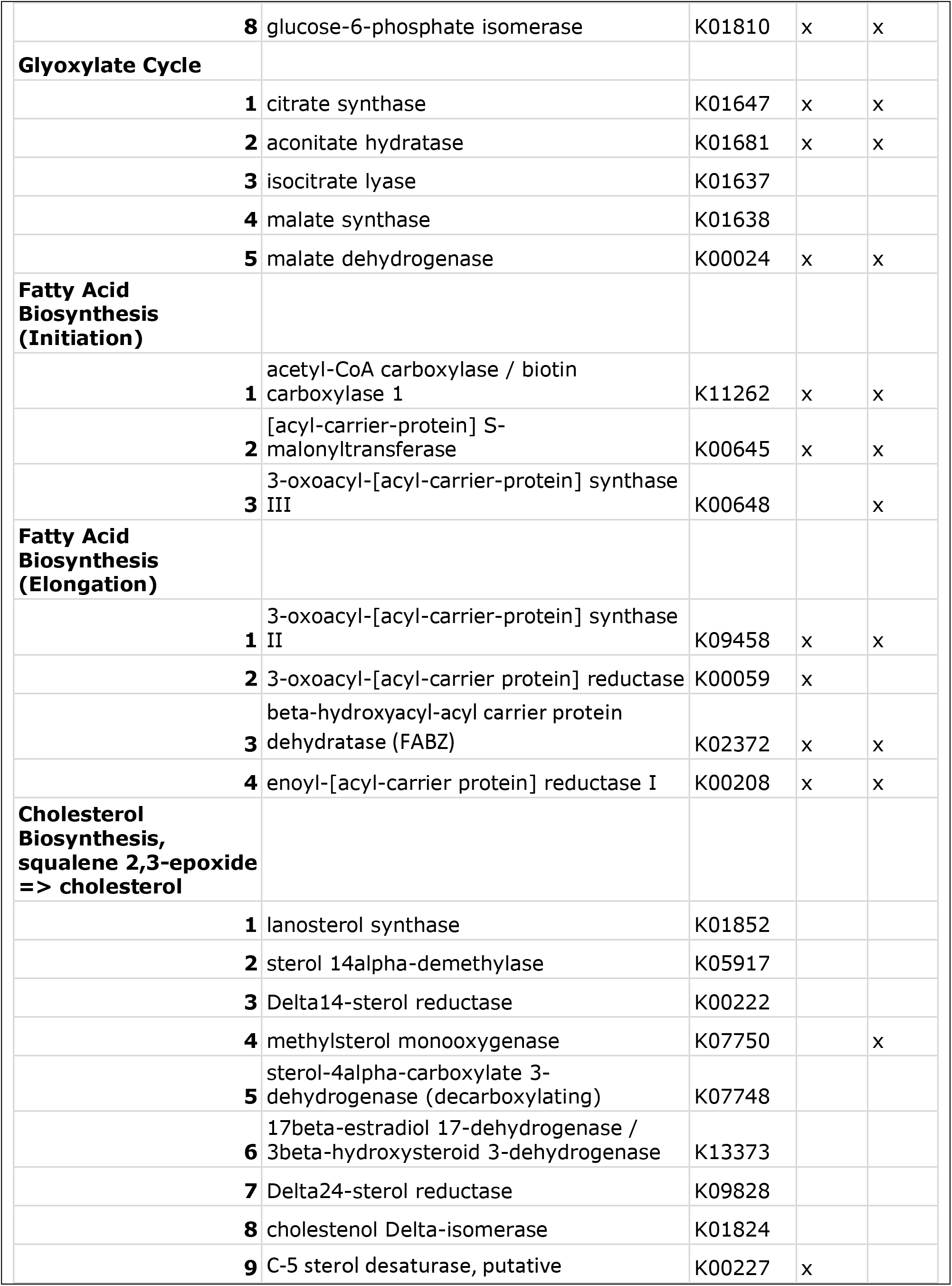
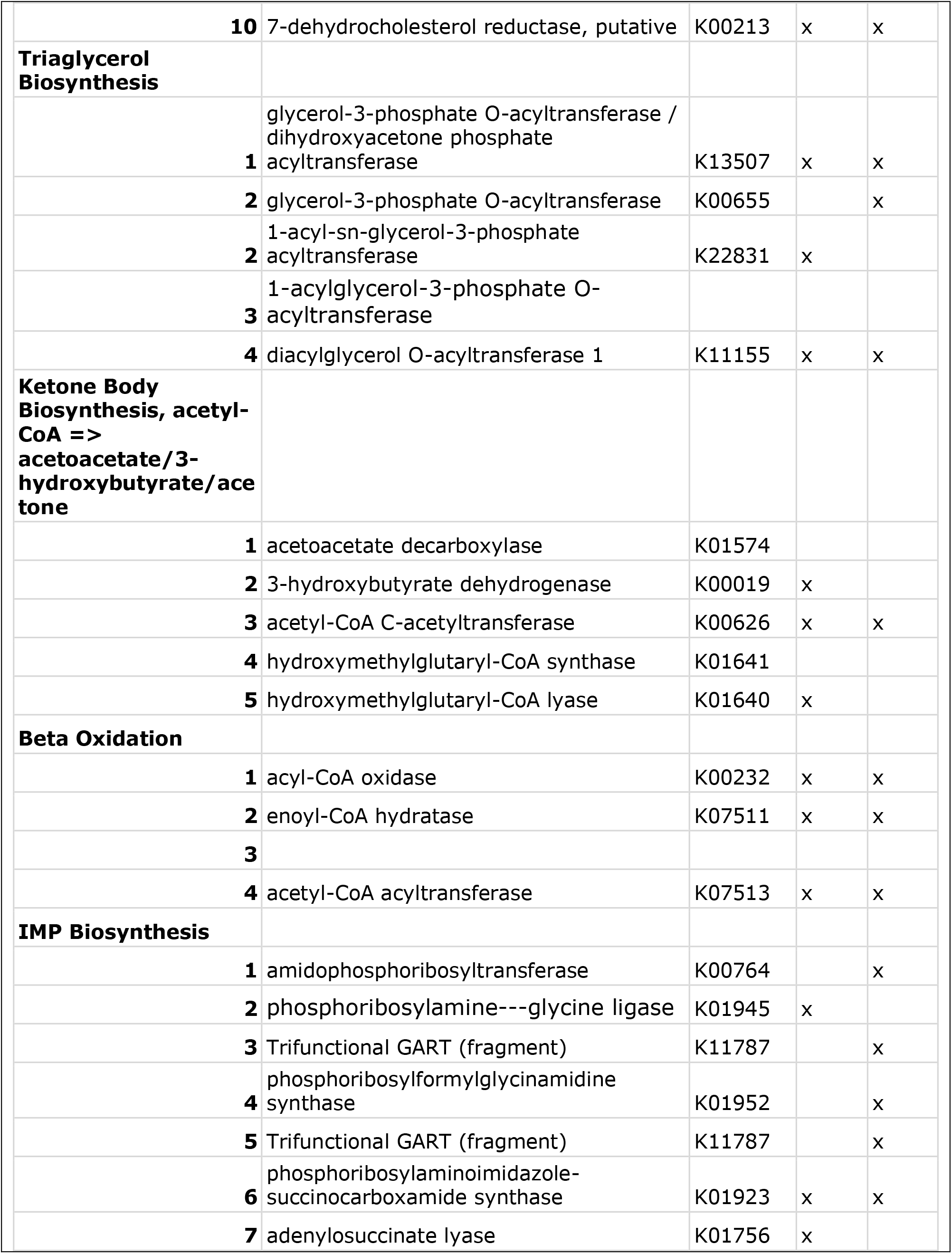
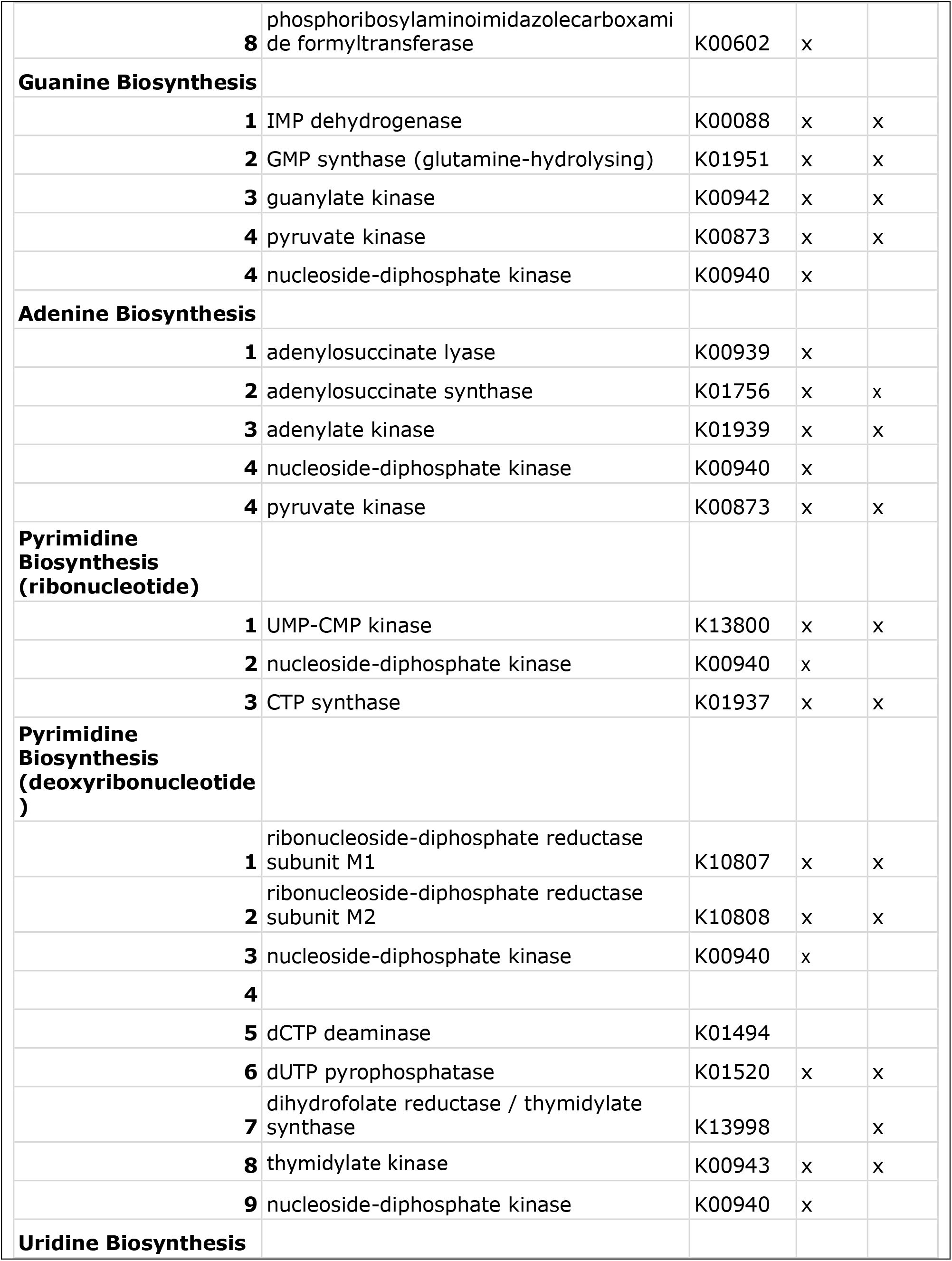
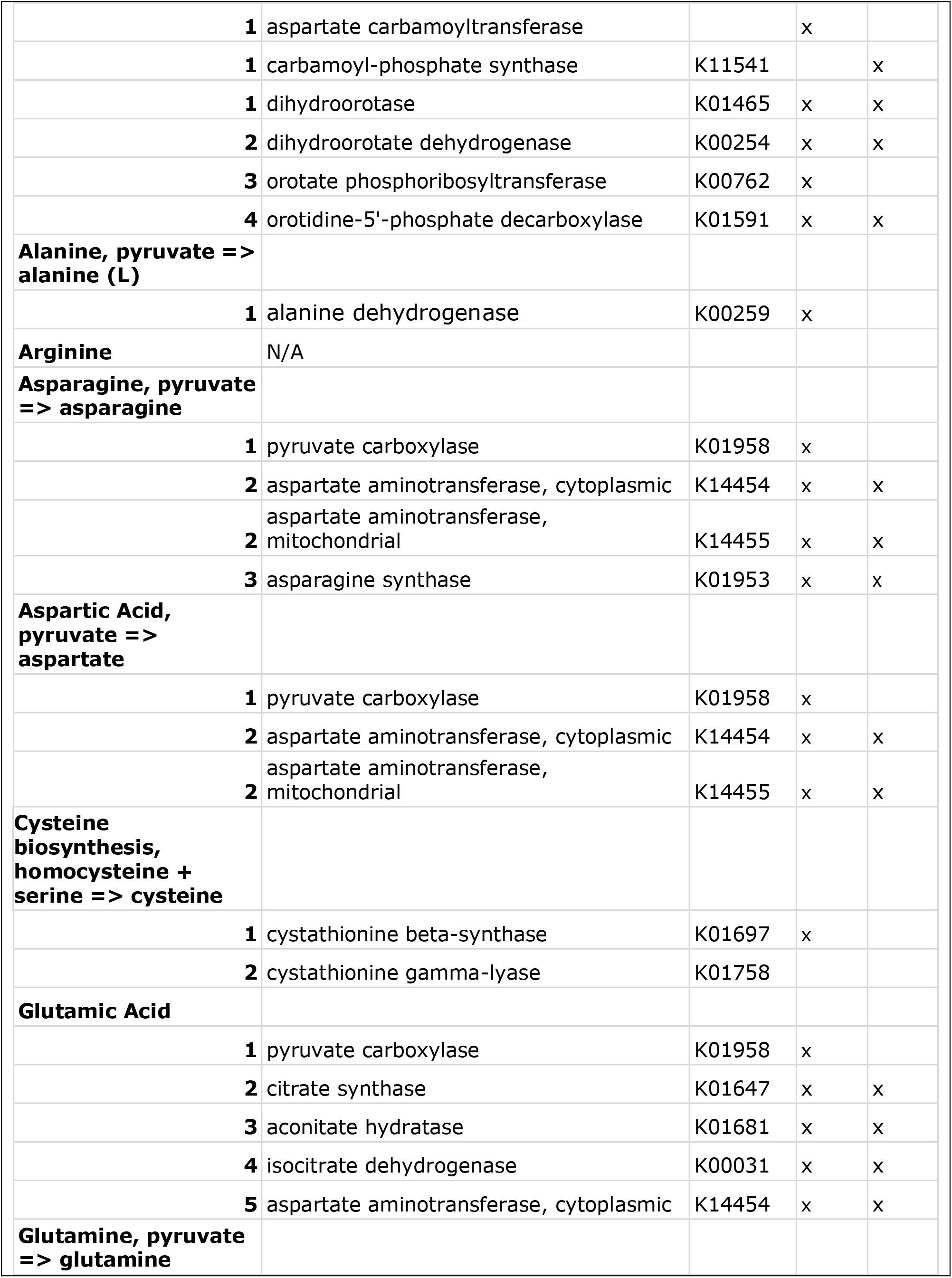
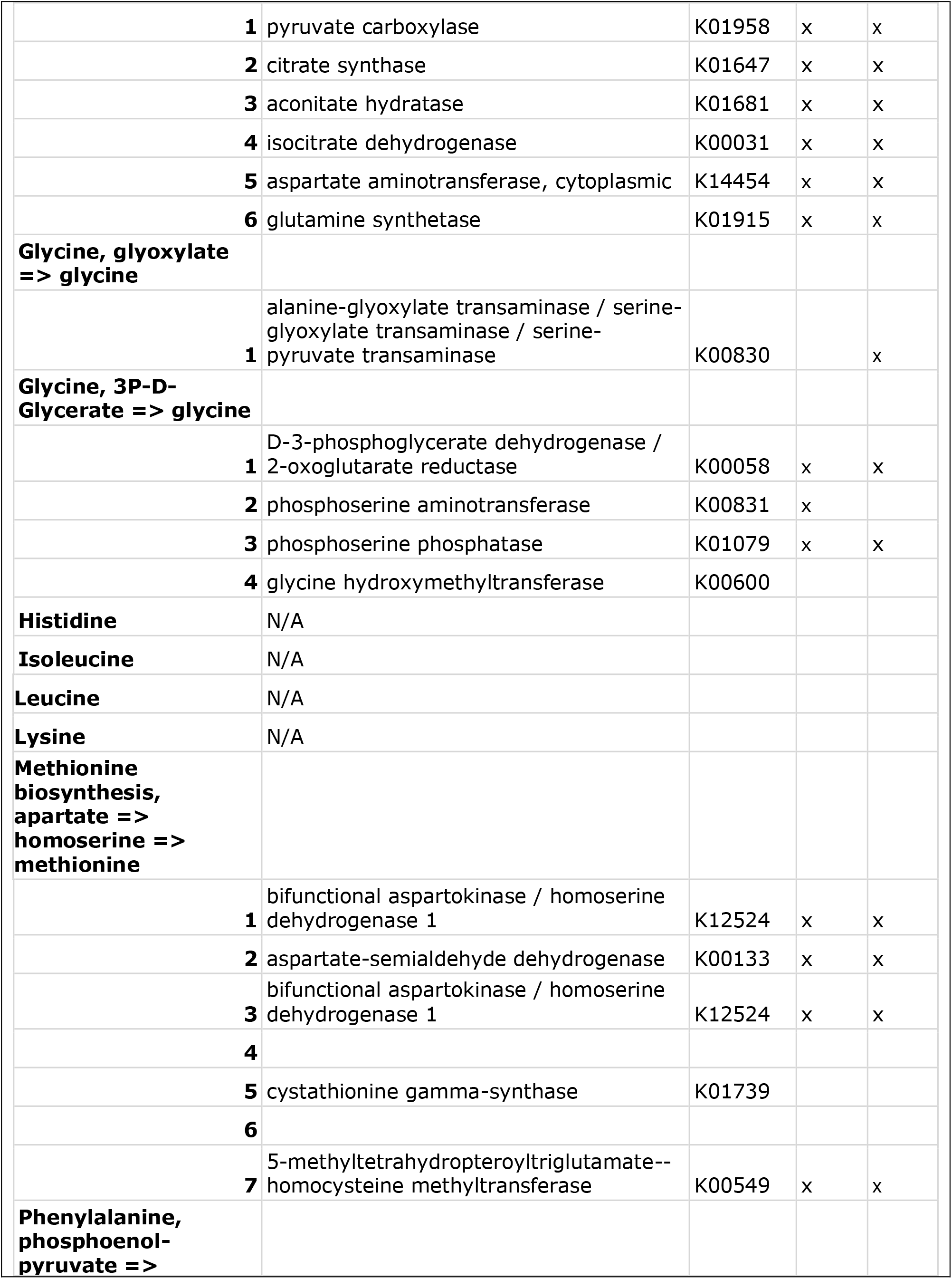
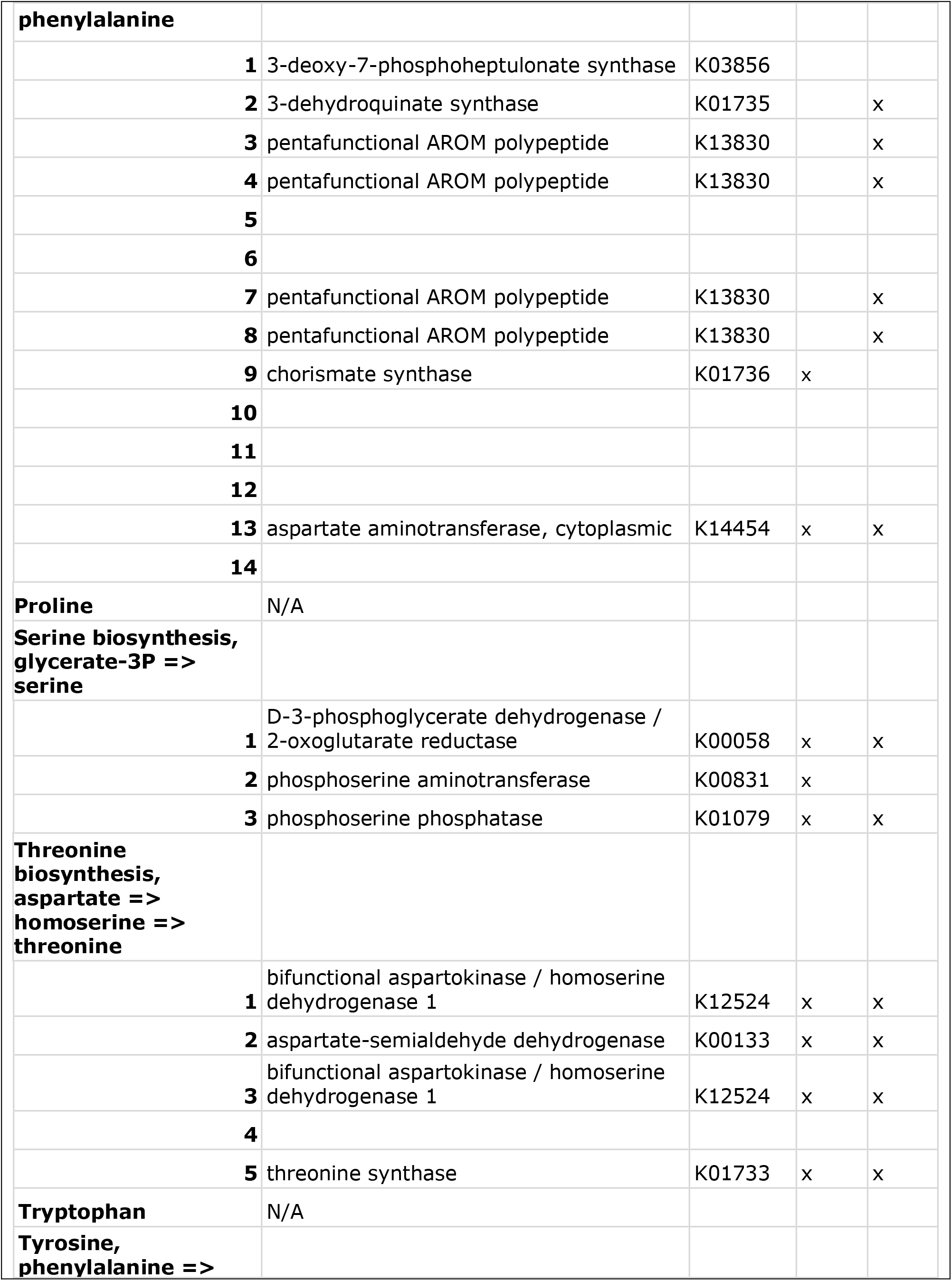
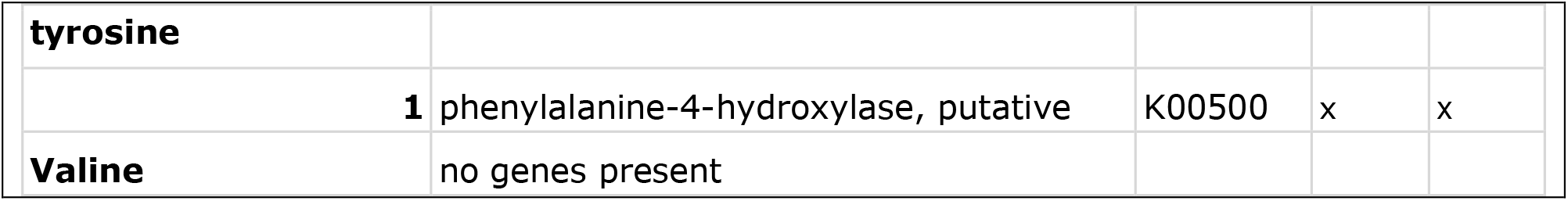
Key functional genes annotations from the genome and transcriptome of *Cardiosporidium cionae*, corresponding to those seen in Figure 2, beginning with the top middle slice and moving clockwise. Annotations are based on functional predictions from KEGG and referenced by the corresponding K-Number.

Supplementary Figure 1: The annotated, circularized *Cardiosporidium cionae* apicoplast. Two *C. cionae* apicoplasts were recovered that were 99.55% similar overall and contained identical gene organization, with the SNPs localized to the sufB gene. The *C. cionae* apicoplasts are similar in size, organization, and gene content to the *Nephromyces* apicoplasts.

Supplementary Figure 2: Comparisons of the bacterial endosymbiont genomes in *Cardiosporidium cionae* (AlphaC) and Nephromyces (AlphaN, Beta, Bac). The left Venn diagram depicts orthologous groups predicted by OrthoFinder, while the right shows functional overlap predicted with KEGG.

