## Supplementary figures and images for "Metabolic contributions of an alphaproteobacterial endosymbiont in the apicomplexan *Cardiosporidium cionae*"

### Supplemental Material

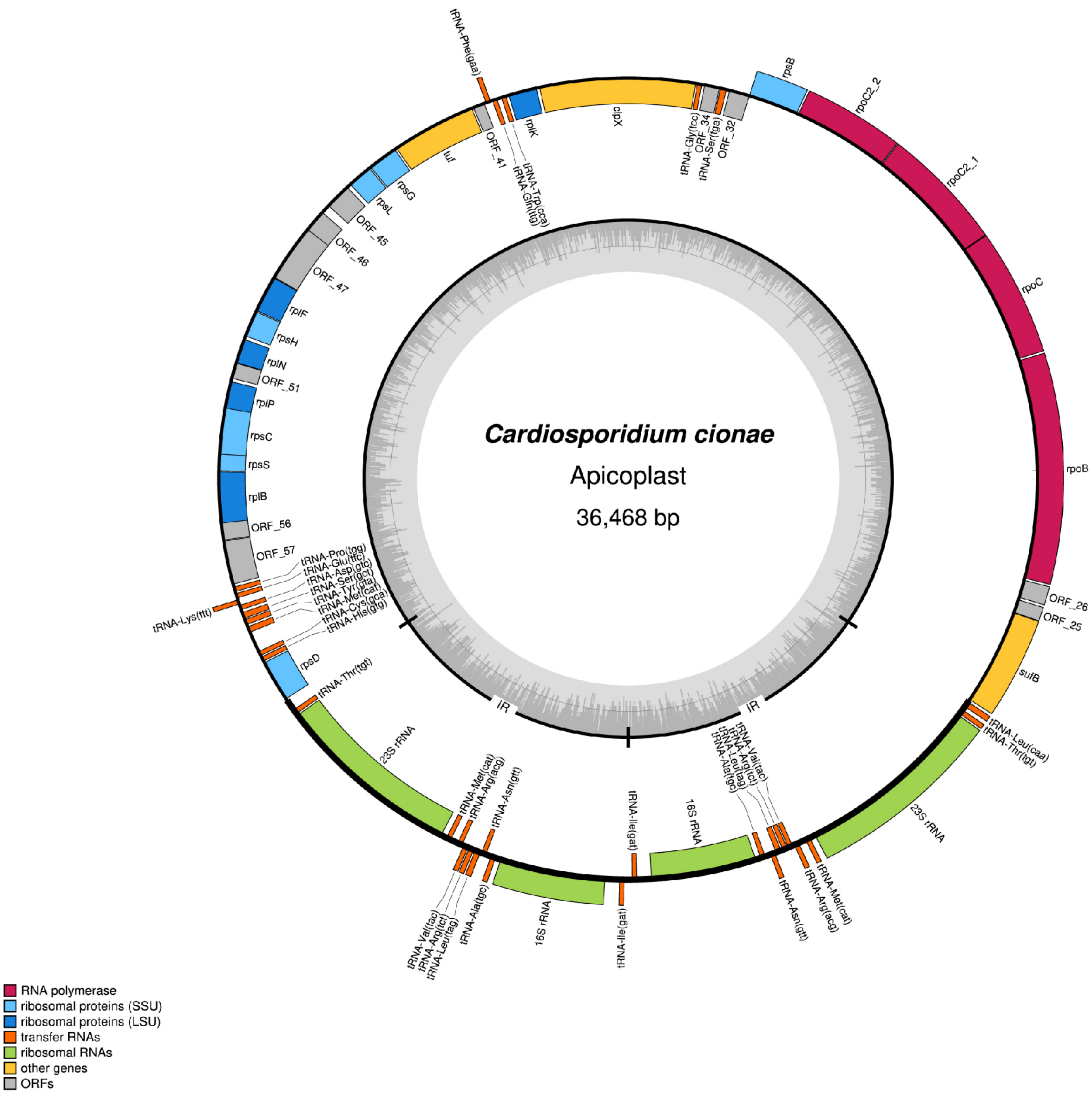

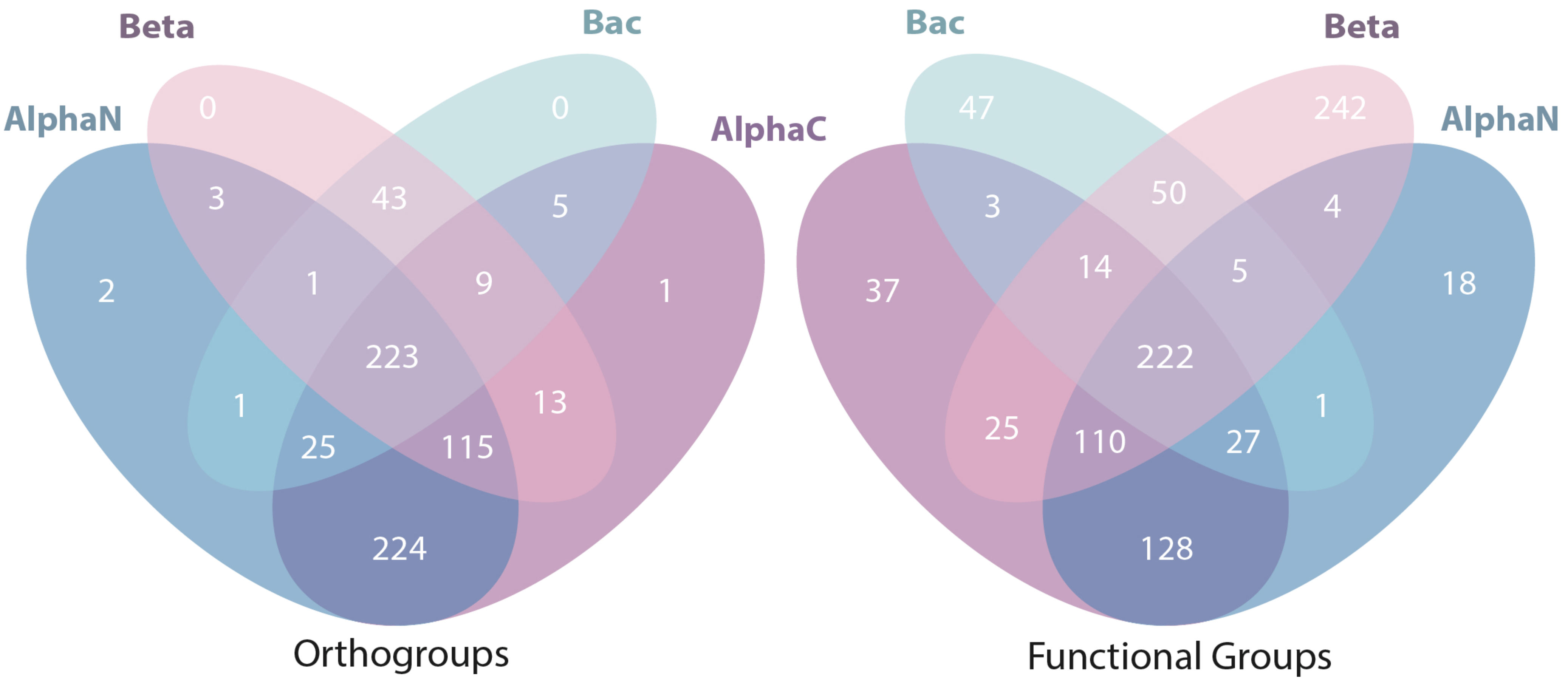

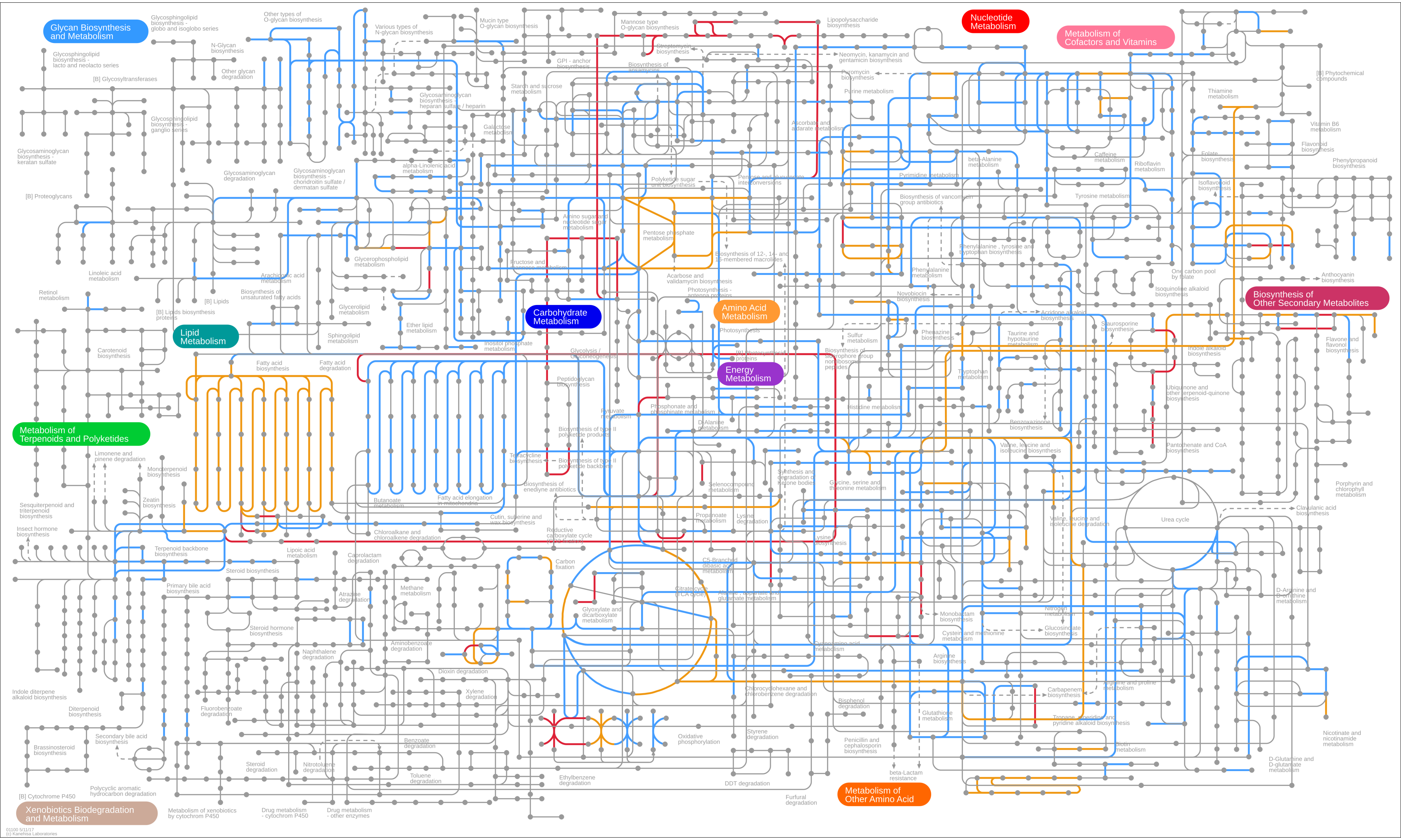
